# Imaging Giant Vesicle Membrane Domains with a Luminescent Europium Tetracycline Complex

**DOI:** 10.1101/2022.07.01.498133

**Authors:** Jennie L. Cawley, Brett A. Berger, Adeyemi T. Odudimu, Aarshi N. Singh, Dane E. Santa, Ariana I. McDarby, Aurelia R. Honerkamp-Smith, Nathan J. Wittenberg

## Abstract

Microdomains in lipid bilayer membranes are routinely imaged using organic fluorophores that preferentially partition into one of the lipid phases, resulting in fluorescence contrast. Here we show that membrane microdomains in giant unilamellar vesicles (GUVs) can be visualized with europium luminescence using a complex of europium (III) and tetracycline (EuTc). EuTc is unlike typical organic lipid probes in that it is a coordination complex with a unique excitation/emission wavelength combination (396/617 nm), a very large Stokes shift (221 nm), and a very narrow emission bandwidth (8 nm). The probe preferentially interacts with liquid disordered domains in GUVs, which results in intensity contrast across the surface of phase-separated GUVs. Interestingly, EuTc also alters GM1 ganglioside partitioning. GM1 typically partitions into liquid ordered domains, but after labeling phase-separated GUVs with EuTc, cholera toxin B-subunit (CTxB), which binds GM1, labels liquid disordered domains. We also demonstrate that EuTc, but not free Eu^3+^ or Tc, significantly reduces lipid diffusion coefficients. Finally, we show that EuTc can be used to label cellular membranes similar to a traditional membrane probe. EuTc may find utility as a membrane imaging probe where its large Stokes shift and sharp emission band would enable multicolor imaging.

## INTRODUCTION

Characterizing lipid membrane spatial heterogeneity is essential to understanding the structure and function of cellular membranes. Cell membranes are asymmetric and may be organized into dynamic, laterally-heterogeneous nanometer-sized membrane domains known as “lipid rafts.” Lipid rafts contain a variety of lipids and proteins involved in cell signaling processes.^1^ Due to the complexity of cell membranes, many researchers turn to model membranes,^2–3^ such as liposomes,^4^ giant unilamellar vesicles (GUVs),^5^ or supported lipid bilayers^6^ (SLBs) for biophysical studies. Model membranes are advantageous because they are able to mimic the properties of cellular membranes^7^ and their composition can be tightly controlled. Depending on their lipid composition, model membranes can possess lipid raft-like liquid ordered (Lo) domains that are enriched in saturated phospholipids, sphingolipids, cholesterol and glycosphingolipids, like gangliosides.^8–9^ A second phase, referred to as the liquid disordered (Ld) domain is enriched in unsaturated lipids. The most common strategy for imaging Lo and Ld domains is fluorescence microscopy, which is typically accomplished by incorporating lipid-conjugated organic fluorophores or polycyclic aromatic hydrocarbons that partition into one phase or the other.^10–11^ Conveniently, these probes are often compatible with “off-the-shelf” filter cubes; however, they have small Stokes shifts and broad emission spectra resulting in fluorescence crosstalk that limits image multiplexing possibilities.

To overcome these limitations and potentially expand the membrane imaging toolbox, we utilize a luminescent complex composed of europium III (Eu^3+^) and tetracycline (Tc) to identify membrane phase heterogeneity among giant unilamellar vesicles (GUVs). Lanthanide luminescence, including Eu^3+^, has been exploited for chemical analysis and biological imaging applications,^12–18^ and here we show that a simple ligand, Tc, enables the imaging of lipid membrane spatial heterogeneity. The luminescence of Eu^3+^ is enhanced by Tc excitation.^19–20^ Tc acts as an antenna, and its excitation is followed by a ligand-to-metal charge transfer, yielding sharp Eu^3+^ emission bands, with the most intense band between 610 and 620 nm. The emission band of EuTc is very narrow in comparison to traditional organic fluorophores. In pure water, the luminescence of EuTc reaches a maximum when the stoichiometry reaches a 1:1 ratio of Eu^3+^:Tc, suggesting a 1:1 stoichiometry of the complex.^19^ The luminescence of the EuTc complex is sensitive to its environment (e.g. pH, hydration shell, other ligands).^19^ Analytes that can enter the Eu^3+^ inner coordination sphere displace water.^21–22^ Coordinated water quenches some EuTc emission by accepting energy from the excited Eu^3+^ states followed by non-radiative decay.^23^ Therefore, displacement of water from the inner coordination sphere causes a significant increase in emission intensity. Due to this phenomenon, the EuTc complex has been applied to the detection of low density lipoprotein (LDL), various surfactants, DNA, hydrogen peroxide, and sialic acid-bearing cancer biomarkers in human plasma.^13, 21, 24–29^ In all cases, the EuTc complex proved to be highly sensitive for the analyte of interest. It also is a suitable alternative to traditional organic fluorescent probes because it is simply prepared, decomposes slowly, has a working pH of ∼7, is fluorescent in buffered systems, and can be used for sensing in dynamic biological systems.^30–31^ Considering these advantages and its unique spectral properties, we suspected that EuTc luminescence is sensitive to phospholipid membranes and can be used for membrane imaging. Here we show that EuTc can be used to visualize membrane heterogeneity in GUVs. We show that EuTc preferentially labels Ld domains, surprisingly causes GM_1_ redistribution into Ld domains, and reduces lipid diffusion coefficients. Finally, we incubated cells with EuTc and observed patterns of fluorescent labeling similar to a traditional organic membrane probe.

## RESULTS AND DISCUSSION

### EuTc Spectral Characteristics and Lipid Sensitivity

To demonstrate that the EuTc complex can be used as a fluorescent probe for lipid membranes, we compared the excitation and emission characteristics of EuTc and Texas Red-DHPE (TR-DHPE), when associated with dioleoylphosphatidylcholine (DOPC) liposomes in 3-morpholinopropane-1-sulfonic acid (MOPS) buffer, pH 7.0. The structures of EuTc, DOPC, and TR-DHPE are shown in Figure 1A. TR-DHPE is a commonly used probe for membrane phase separation that partitions into the Ld phase.^11^ As shown in Figure 1B, the excitation maximum for EuTc and TR-DHPE are 396 nm, and 589 nm, respectively. There are two major emission peaks of the EuTc complex centered at 592 nm and 617 nm, which correspond to the ^5^D_0_→^7^F_1_ and ^5^D_0_→^7^F_2_ transitions of Eu^3+^, respectively.^32^ The intense, sharp EuTc emission peak at 617 nm has a very narrow bandwidth, especially when compared to traditional organic fluorophores. Specifically, the 617 nm emission peak of EuTc has a full width at half maximum (FWHM) of 8 nm, and it is red-shifted 221 nm from the excitation maximum. For comparison, the TR-DHPE emission peak has a FWHM of 32 nm with a Stokes shift of only 20 nm. Figure 1C displays that in the absence of the ligand, Tc, Eu^3+^ luminescence is negligible in the presence of DOPC liposomes. This is expected as free Eu^3+^ has a small molar absorptivity (ε < 1 M^−1^ cm^−1^)^33^ because the pertinent electronic transitions are forbidden.^32^

**Figure 1.**
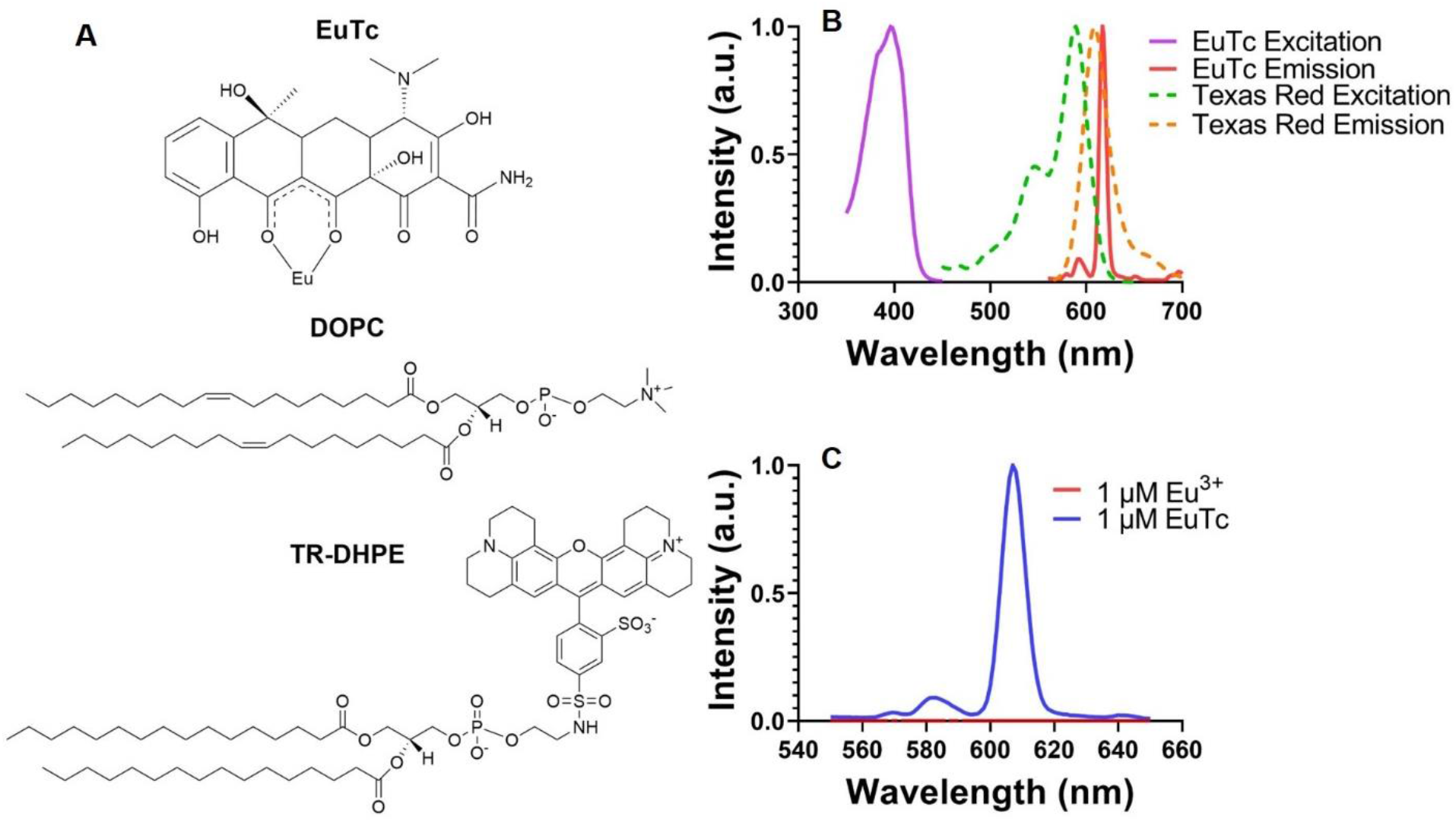
Molecular structures and spectroscopic characteristics of EuTc (1 μM) and Texas Red-DHPE with DOPC liposomes. (A) Molecular structures of EuTc, DOPC, and TR-DHPE. (B) Excitation and emission of spectra of EuTc compared with excitation and emission spectra of Texas-Red DHPE. (C) Emission spectra of the EuTc complex (1 μM) and Eu^3+^ (1 μM) alone in the presence of DOPC liposomes. EuTc and Eu^3+^ were excited at 400 nm.

Next, we sought to determine if EuTc emission is enhanced by phospholipids in lamellar lipid structures. Previous research has shown that various surfactants (sodium dodecylsulfate (SDS), cetylpyridinium chloride (CPC), Triton-X 100, Brij 58 and Brij 78), all which form micellar structures, can enhance the emission of EuTc.^27^ First, we formed liposomes by a series of freeze-thaw cycles and extrusion through a 100-nm pore filter. The liposomes were then exposed to 1 μM EuTc. The liposomes were composed of DOPC, and the lipid concentration in the liposome suspensions ranged from 10 nM to 100 μM, thus the molar ratio of DOPC to EuTC ranged from 1:100 to 100:1. EuTc emission depended on total lipid concentration, and increased with increasing lipid concentration (Fig. 2), with an emission enhancement of greater than 5-fold when 100 μM DOPC was present. This confirms that EuTc emission, while observed in the absence of lipids, is sensitive to the concentration of lipids in lamellar structures in an aqueous environment. In earlier work that examined surfactant-enhanced EuTc luminescence, molar ratios of surfactant to EuTc ranged from 1550:1 to 3470:1.^27^ With these ratios, EuTc luminescence was enhanced from 7.6 to 30.3-fold, depending on the surfactant. Assuming these enhancements scale with surfactant to EuTc molar ratio, comparing them to the enhancement we observed with 100:1 ratio of DOPC to EuTc shows that the DOPC liposomes result in significantly greater EuTc emission enhancement. The luminescence of the EuTc complex is known to increase upon displacement of water molecules from the coordination sphere. Our results suggest that EuTc interacts with the lipid bilayer membrane in a manner that diminishes non-radiative decay pathways, similar to that of water displacement.

**Figure 2.**
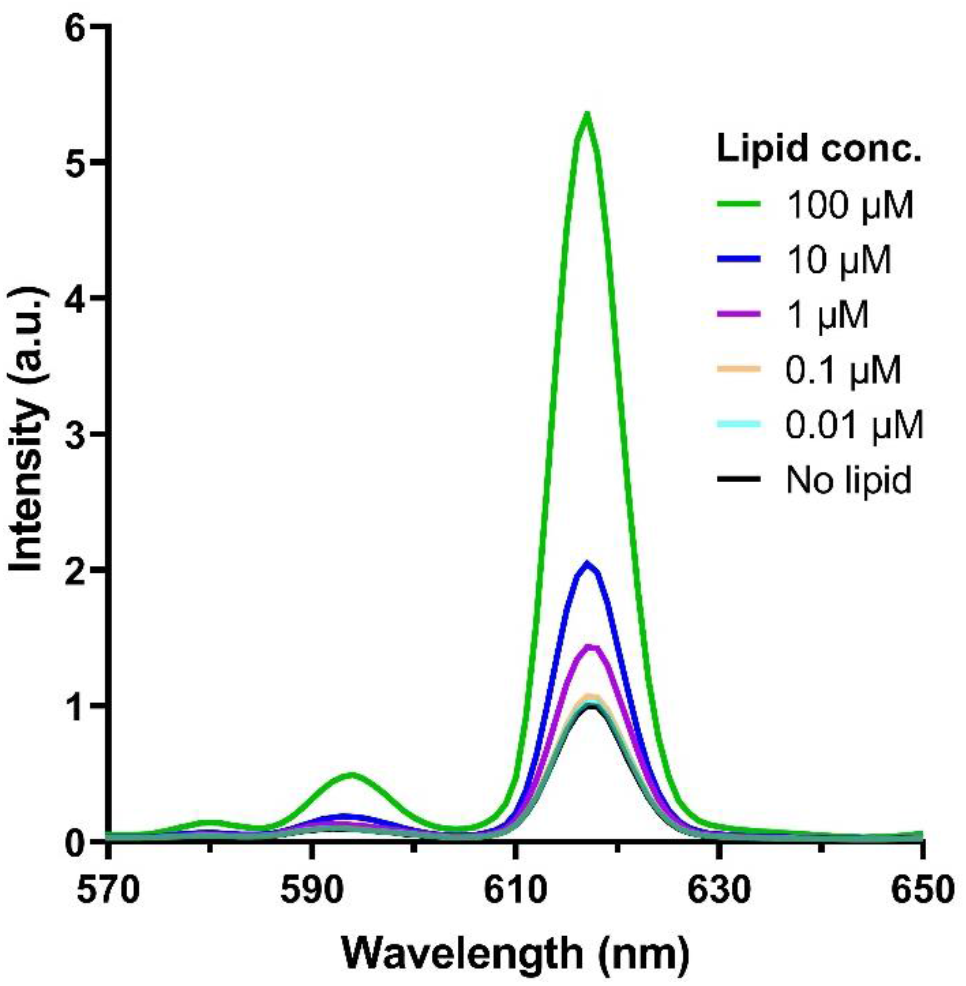
EuTc emission intensity in the presence of varying concentration of DOPC liposomes. The DOPC liposomes enhance EuTc emission intensity in a concentration-dependent manner.

### GUV Imaging with EuTc

In addition to showing spectroscopically that EuTc luminescence is enhanced in the presence of lipids, we sought to demonstrate how the EuTc complex could be used as an imaging probe. To demonstrate this, we prepared single phase (DOPC) and phase separating GUVs containing a mixture of DOPC, dipalmitoylphosphatidylcholine (DPPC), and cholesterol (DOPC/DPPC/cholesterol; 40:40:20 molar ratio) and imaged them in the presence of 1 μM EuTc. The structures of cholesterol and DPPC are shown in Figure 3A. Other than EuTc, no other labels were present. For EuTc imaging, a custom filter set was employed (Fig. S1). Labeling DOPC GUVs with EuTc resulted in uniform membrane luminescence (Fig. 3B). In contrast, phase separating GUVs displayed EuTc luminescence over only a portion of the GUV membrane (Fig. 3C). A hallmark of a membrane phase sensitive probe is that it labels one membrane phase more intensely than the other. Based on our observations here, we conclude that EuTc is a phase sensitive probe.

**Figure 3.**
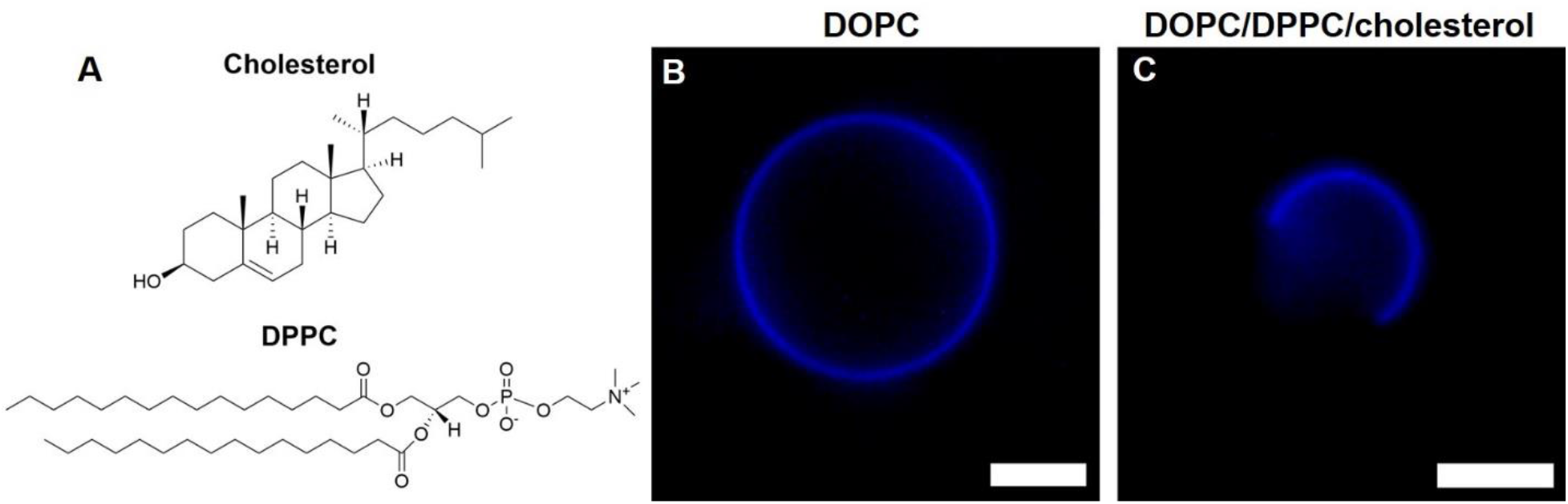
EuTc labeled GUVs. (A) Molecular structurs of DPPC and cholesterol. (B) DOPC GUV and (C) phase separated GUV (DOPC/DPPC/cholesterol, 40:40:20) labeled with 1 µM EuTc. All scale bars are 5 µm.

To further explore EuTc for vesicle imaging, we prepared additional single phase GUVs with three different lipid compositions: DOPC/cholesterol (80:20), DOPC/GM1 (98:2), and DOPC/GM1/cholesterol (78:2:20).^7, 34^ GUVs were all presumably in a single Ld phase and were imaged in the presence of 1 μM EuTc, and uniform EuTc luminescence on the membrane was observed (Fig. 4A). In addition to the EuTc complex, a fluorescent conjugate of cholera toxin subunit-B (CTxB-FITC) was added to each imaging chamber containing the GUVs. CTxB-FITC binds GM1 with high affinity^35^ and its fluorescence was only observed with GUVs composed of DOPC/GM1 (98:2) and DOPC/GM1/cholesterol (78:2:20) (Fig. 4B). An overlay of EuTc and CTxB-FITC channels display colocalization on the GUV membranes only for GUVs possessing GM1 (Fig. 4C). Fluorescence intensity profiles along the dashed lines in Fig. 4 are shown in Fig. S2. Our observations confirm that EuTc can be used to visualize membranes containing GM1 and/or cholesterol and the presence of the EuTc probe does not appear to significantly alter CTxB-FITC binding to GM1. Interestingly, free Eu^3+^ has been shown to inhibit the binding of complete cholera toxin (A and B subunits) and amyloid-beta oligomers to lipid membranes containing GM1.^36–37^ However, we see no evidence that EuTc acts in a similar manner.

**Figure 4.**
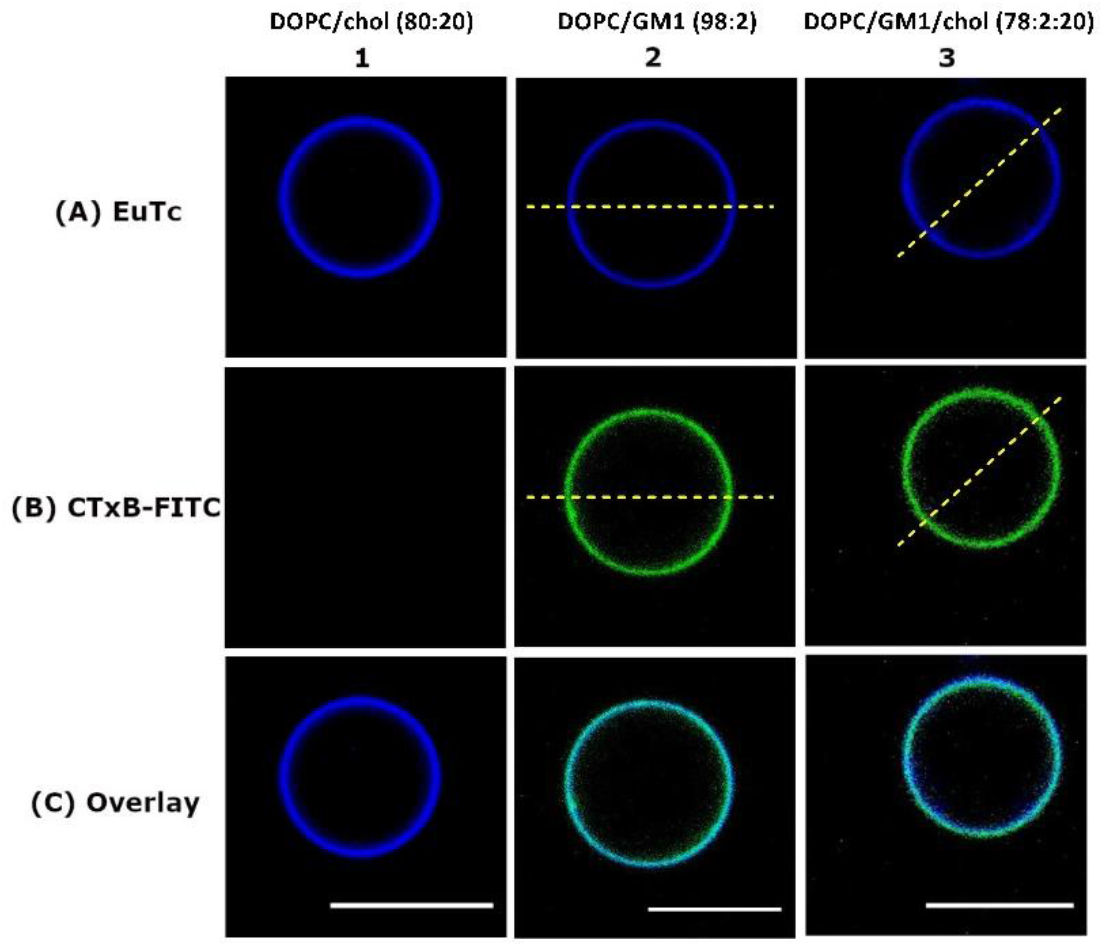
Single phase GUVs labeled with EuTc and CTxB-FITC. The GUV compositions are noted above the column numbers. Rows indicate fluorescence due to (A) EuTc and (B) CTxB-FITC labeling. Overlay images (C) show colocalization of EuTc and CTxB-FITC on the GUV membrane only when GM1 is present. Intensity profiles along the yellow dashed lines in Columns 2 and 3 can are shown in Fig. S2. All scale bars are 10 µm.

As an additional control, single phase GUVs composed of DOPC/GM1/TR-DHPE (98:1:1) were prepared and labeled with CTxB-FITC. The TR-DHPE and CTxB-FITC signals colocalize on the GUV membrane, as expected (Fig. S3). It is important to note that EuTc was not washed out of the chamber prior to collecting the images shown in Figs. 3 and 4. Despite this, the images have good signal to noise ratios due to the significant luminescence enhancement of EuTc upon its interaction with lipid bilayer membranes.

The intensity contrast of EuTc labeling on phase separated GUVs (Fig. 3C) could be explained by two different scenarios. In one scenario, EuTc may preferentially localize with one membrane phase or the other (Ld or Lo). This would increase its local concentration and thus increase the luminescence intensity of that region. Alternatively, EuTc may localize equally to both phases (i.e. equal concentration across both phases), but become more luminescent when interacting with one phase or the other. However, these two scenarios are not mutually exclusive and some combination of them may be responsible for the intensity contrast.

### EuTc Colocalizes with a Ld Domain Marker

Thus far, we have identified that the EuTc complex can be used as a fluorescent imaging probe for single phase GUVs and labels phase-separated membranes heterogeneously. In an attempt to determine which phase of the GUV (Ld or Lo) the EuTc probe is labeling, we prepared a series of GUVs with lipid compositions chosen to lie along a tie line in the DOPC-DPPC-cholesterol phase diagram.^38^ A phase diagram and a discussion of tie lines can be found in the Supporting Information (Fig. S4). These compositions yielded vesicles containing different area fractions of Lo and Ld phases with the same composition (Fig. 5A). TR-DHPE was included to mark the Ld domains. Upon incubation with EuTc, we observed that the area fraction labeled with EuTc enlarged as the area fraction of the Ld phase increased (Fig. 5B). Finally, the TR-DHPE and EuTc signals colocalize (Fig. 5C). Fluorescence intensity profiles taken across the vesicles in Fig. 5 are shown in Fig. S5. While the relative intensities of EuTc and TR-DHPE vary from vesicle-to-vesicle, the linescans further demonstrate colocalization of EuTc and TR-DHPE probes. Taken together, this suggests that EuTc labels Ld domains.

**Figure 5.**
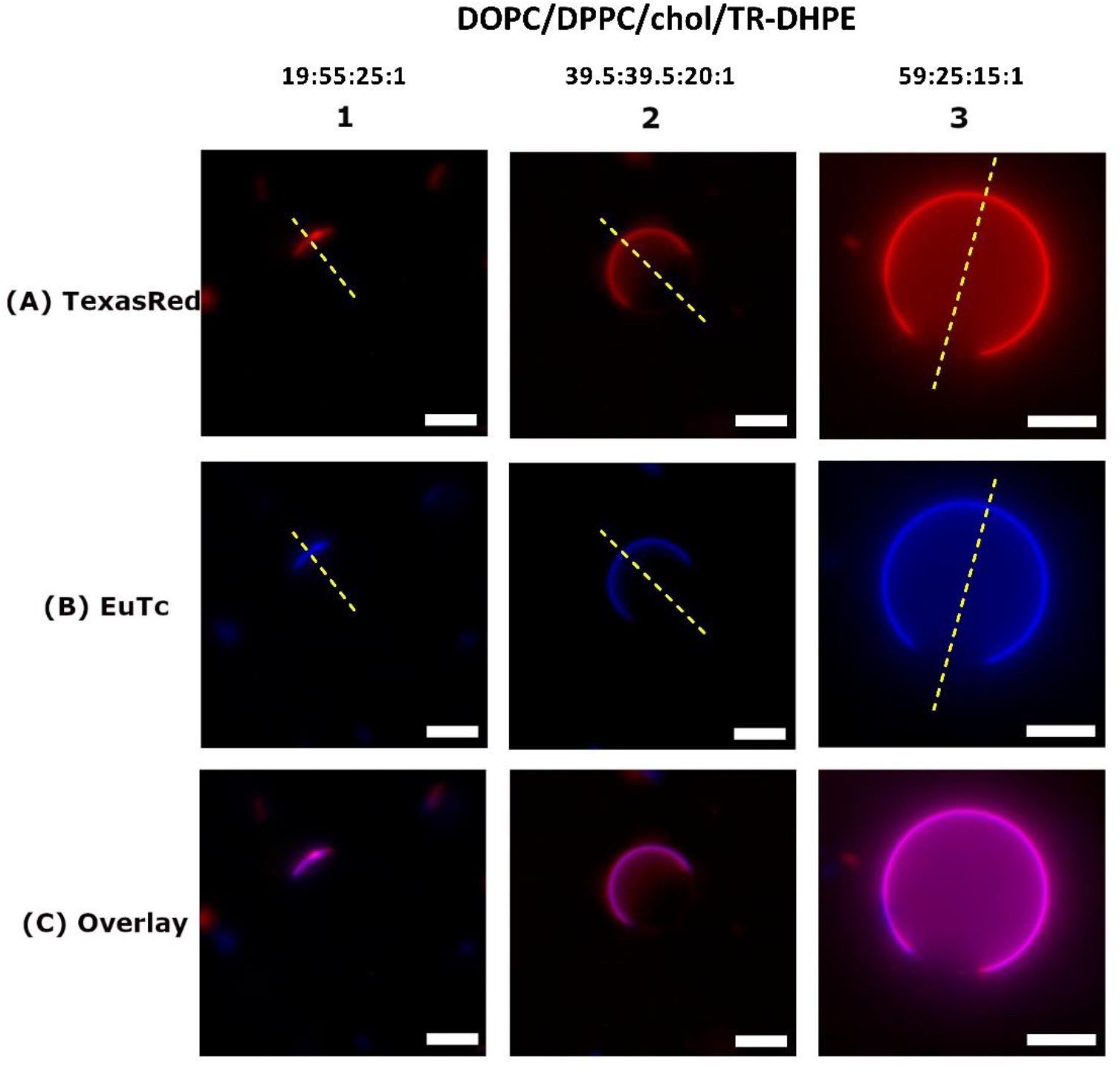
Phase separating GUVs composed of DOPC, DPPC, cholesterol, and TR-DHPE with varying area fraction of the Ld and Lo phases imaged by Texas Red (TR-DHPE) and EuTc luminescence. The ratios above the column numbers indicate the molar ratios of DOPC, DPPC, cholesterol, and TR-DHPE in the GUV. Rows indicate fluorescence due to (A) Texas Red-DHPE and (B) EuTc labeling. Overlay images (C) show colocalization of EuTc and Texas Red on the GUV membrane. Fluorescence intensity profiles along the yellow dashed lines are shown in Fig. S5. All scale bars are 5 µm.

### EuTc Redistributes GM1

To further explore EuTc labeling of Ld domains, we sought to concurrently label Ld and Lo domains. In this set of experiments, we used CTxB binding to GM1 to label the Lo domains. GM1 is known to strongly partition into Lo domains in phase separated GUVs,^39^ thus binding of GM1 by fluorescent CTxB will indicate the position of the Lo domains. We prepared GUVs composed of DOPC/DPPC/cholesterol/GM1 (39.5:39.5:20:1) and labeled the GUVs first with 10 nM CTxB-Alexa647 and then with 1 μM EuTc. Based on prior precedent, we presume CTxB-Alexa647 will label the Lo phase. Surprisingly, Figure 6, column 1 shows that the EuTc labels the same region of the GUV as CTxB-Alexa647. This is an unexpected result because when we prepared phase separating GUVs and labeled them with TR-DHPE, a Ld label, TR-DHPE and EuTc colocalized (Fig 5; Fig. 6, column 4). To evaluate this further, we prepared GUVs in a way in which all labels would be present. These GUVs consisted of DOPC/DPPC/cholesterol/GM1/TR-DHPE, and were exposed to the CTxB-Alexa647 and EuTc labels in a stepwise fashion. The GUVs were first exposed to CTxB-Alexa647, imaged, and then subsequently exposed to EuTc and imaged again. Upon the addition of CTxB-Alexa647, we observed that CTxB-Alexa647 and TR-DHPE label oppositely to one another as expected (Figure 6, column 2). It is well established that GM1 partitions into the Lo phase, thus CTxB-Alexa647 is a Lo phase label.^39^ Conversely, TR-DHPE is a Ld phase label.^11^ Next, we exposed the GUVs to the EuTc complex. Upon imaging the GUVs, we observed that all three labels, EuTc, CTxB-Alexa647, and TR-DHPE colocalize (Figure 6, column 3). The persistent colocalization of the EuTc and TR-DHPE labels confirm that the CTxB-Alexa647 does not impede TR-DHPE partitioning or EuTc labeling. A fluorescence intensity profile along the dashed line in Figure 6, column 3 is shown in Figure S6.

**Figure 6.**
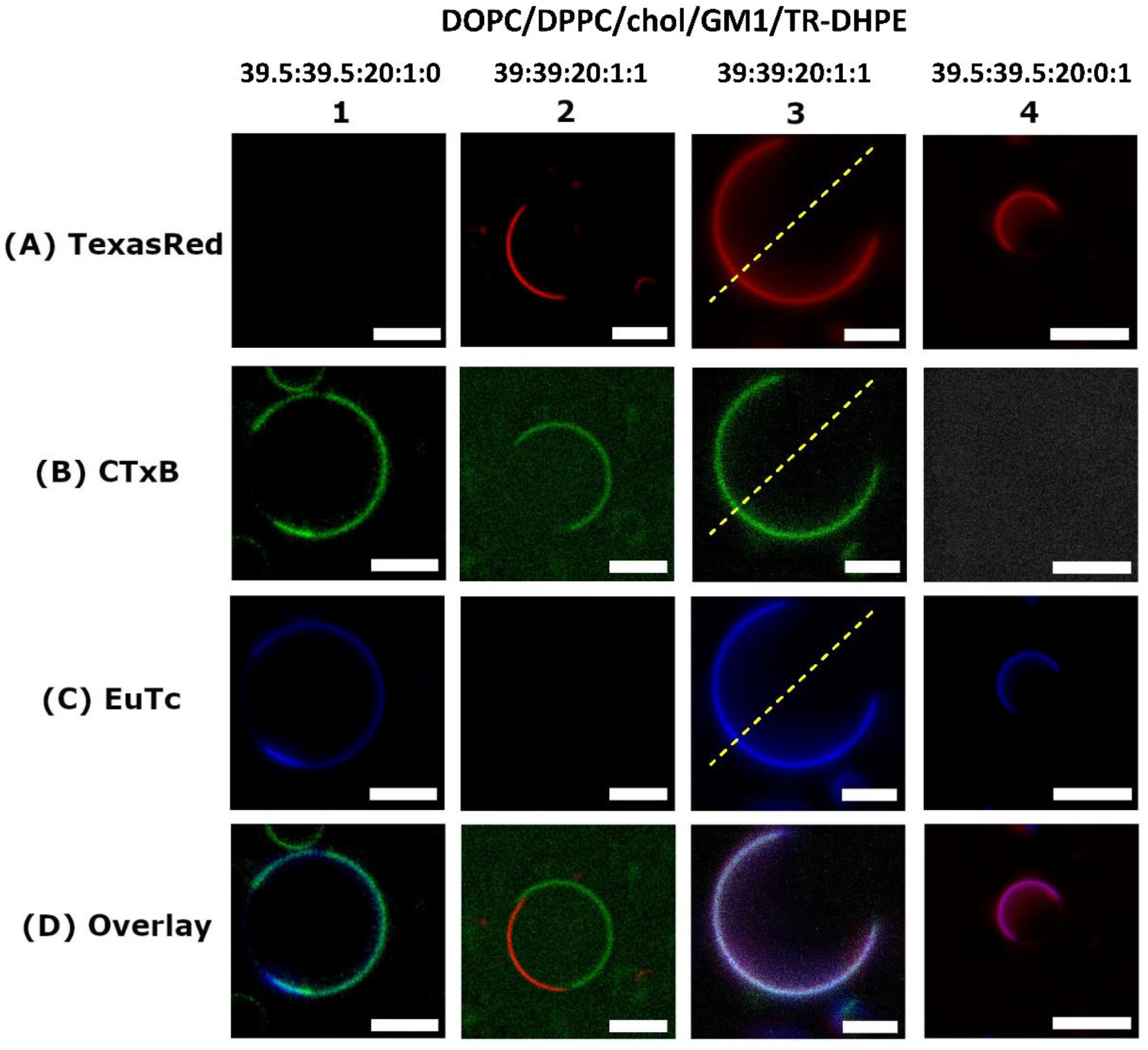
Imaging phase separation in GUVs. Phase separating GUVs imaged by either Texas Red (TR-DHPE) and/or EuTc luminescence and/or CTxB-AlexaFluor647. The GUVs were composed of DOPC, DPPC, cholesterol, GM1, and TR-DHPE. The ratios above the column labels are the molar ratios of the lipids in the GUVs. Rows indicate fluorescence due to (A) Texas Red-DHPE, (B) CTxB-AlexaFluor647, and (C) EuTc labeling. Overlay images (D) colocalization of the various probes. Images shown depict the GUVs in their final state, i.e. with all labels present for a given condition. Fluorescence intensity profiles along the dashed lines are shown in Fig. S6. All scale bars are 5 µm.

It is possible that the labeling order (CTxB-Alexa647 first, EuTc second) may influence our observations. To determine if this is the case, we labeled DOPC/DPPC/cholesterol/GM1 (39.5:39.5:20:1) GUVs first with EuTc, followed by CTxB-Alexa647. We observed that EuTc and CTxB-Alexa647 again colocalize (Fig. 7). In summary, we observe that regardless of labeling order, the EuTc colocalizes with CTxB-Alexa647. Additionally, the CTxB-Alexa647 intensity intensities before and after addition of EuTc are similar (Fig. S7). Furthermore, when all labels (TR-DHPE, CTxB-Alexa647, and EuTc) are present, all labels colocalize. However, when EuTc is not present, TR-DHPE and CTxB-Alexa647 do not colocalize. This suggests that EuTc alters the phase preference of GM1 or the GM1/CTxB complex, though the underlying causes of this change remains to be determined.

**Figure 7.**
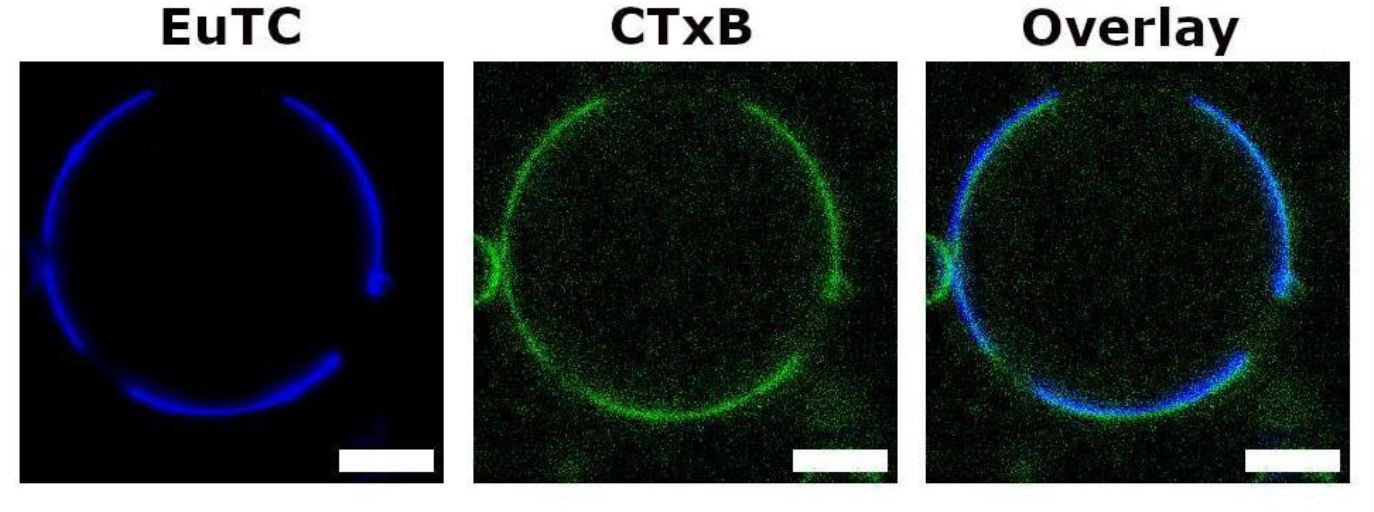
DOPC/DPPC/cholesterol/GM1 (39.5:39.5:20:1) GUVs were labeled with EuTc and then with CTxB-Alexa647. Labeling order (CTxB-Alexa647 then EuTc vs EuTc then CTxB-Alexa647) does not influence colocalization. All scale bars are 5 µm.

### EuTc reduces lipid diffusion coefficients

To examine if EuTc alters the physicochemical properties of membranes, we used fluorescence recovery after photobleaching (FRAP) to determine lipid diffusion coefficients. We prepared supported lipid bilayers (SLBs) consisting of DOPC/TR-DHPE (99:1) and DOPC/TopFlourPC (99:1), and DOPC/BODIPY-GM1 (99:1). The structures of TopFluorPC and BODIPY-GM1 are shown in Figure 8A. After SLB formation, FRAP was conducted before and after adding EuTc (1 μM) to the aqueous medium. As shown in Figure 8B-D, with all compositions there is an obvious reduction in the rate of fluorescence recovery after EuTc incubation. Fluorescence images of the FRAP recovery process are shown in Fig. S8. The reduction in recovery rates translates to a significant reduction in lipid diffusion coefficients after EuTc exposure (Fig. 9). In the absence of EuTc, all of the fluorophores have diffusion coefficients in the range of 1-2 μm^2^/s, which is indicative of freely diffusing molecules on a solid support.^40^ With DOPC/TR-DHPE, the diffusion coefficient drops from 1.67 ± 0.11 μm^2^/s to 0.55 ± 0.12 μm^2^/s (mean ± S.D.) after EuTc exposure, while with DOPC/TopFluorPC, the diffusion coefficient is reduced from 1.98 ± 0.08 μm^2^/s to 0.59 ± 0.08 μm^2^/s (Fig. 9). We chose the TR-DHPE and TopFluorPC probes for this experiment because their fluorescent moieties are linked to different parts of the phospholipid. The TR group of TR-DHPE is linked to the polar head group and exposed to the aqueous environment, and therefore may have more interaction with EuTc. On the other hand, the TopFluor group of TopFluorPC is linked to the hydrophobic tail. Lipid tail-linked TopFluor has been shown to reside deeply within the hydrophobic core of the bilayer.[30] Additionally, TopFluorPC presents the same phosphocholine headgroup as the background lipid DOPC.

**Figure 8.**
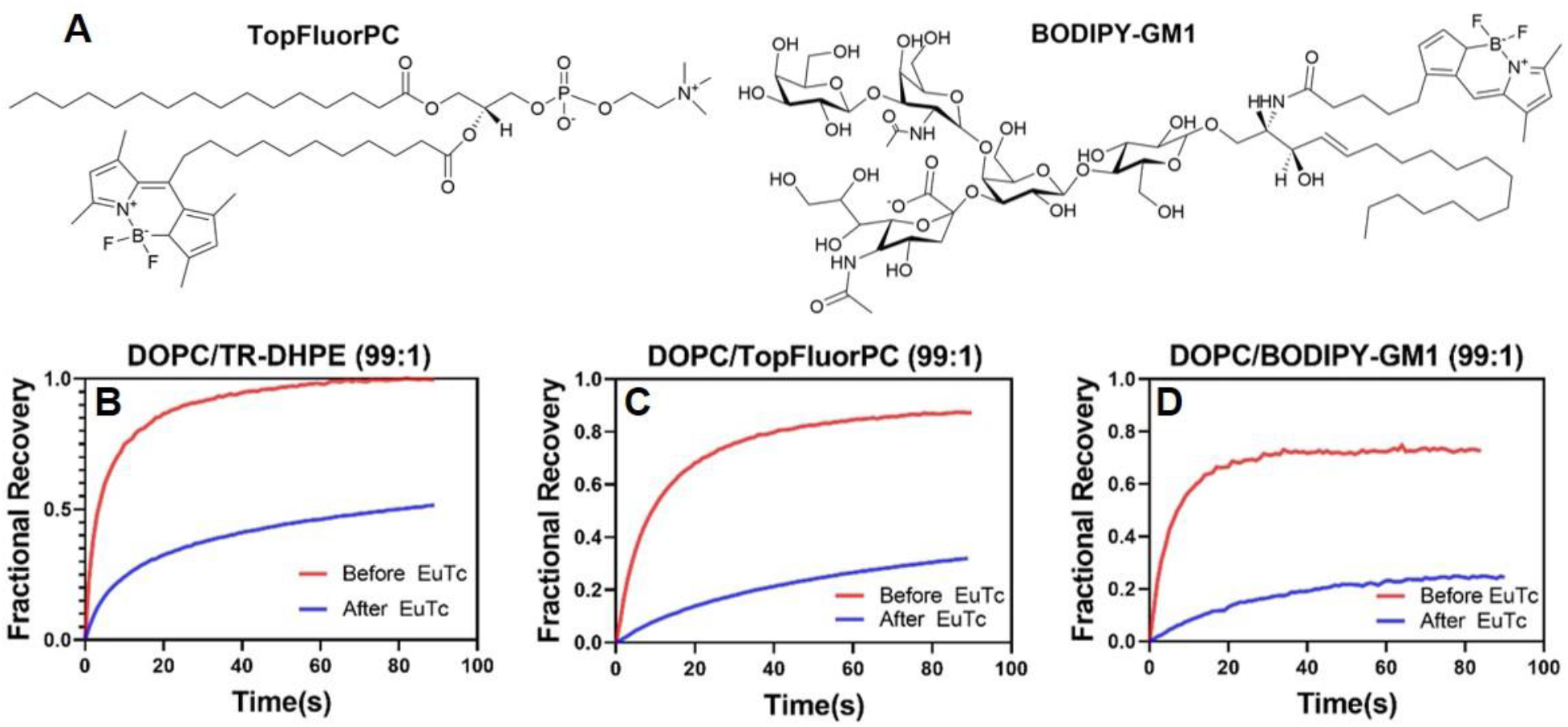
Molecular structures of TopFluorPC and BODIPY-GM1 (A). FRAP recovery curves of DOPC/TR-DHPE (99:1) (B), DOPC/TopFlourPC (99:1) (C), and DOPC/BODIPY-GM1 (99:1) (D) SLBs before and after EuTc exposure.

**Figure 9.**
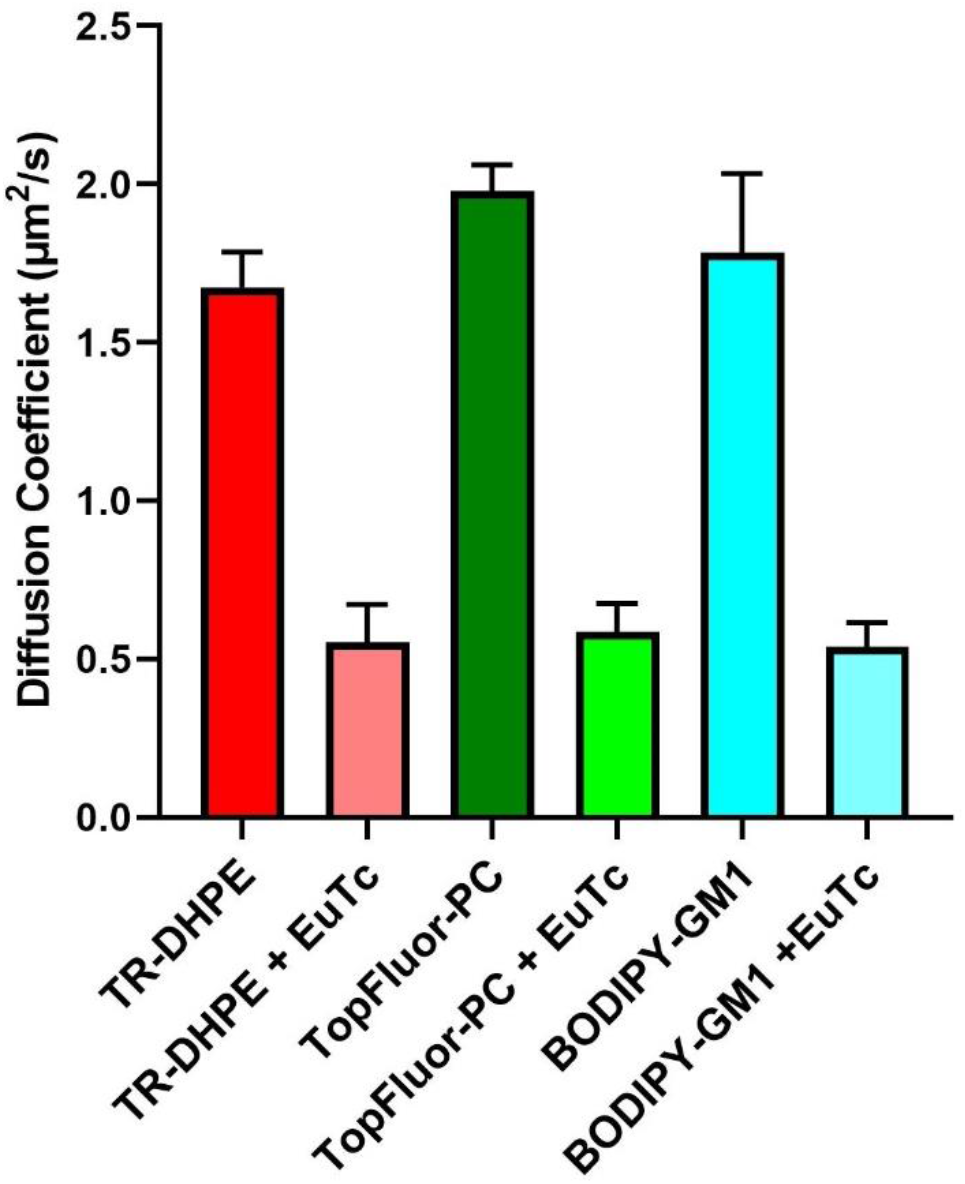
Influence of EuTc on lipid diffusion coefficients in SLBs. Diffusion coefficients of TR-DHPE, TopFluorPC, and BODIPY-GM1 in DOPC SLBs before and after addition of EuTc (+ EuTc). Data is represented as mean ± standard deviation, N ≥ 9 for all samples.

Next, we wanted to determine if EuTc had an effect on the diffusion of GM1. To evaluate this, we utilized a fluorescent analog of GM1, BODIPY-GM1, and interrogated its diffusion in DOPC bilayers using FRAP. In SLBs consisting of DOPC/BODIPY-GM1 (99:1) we observed a similar reduction in recovery rates (Fig. 8D) and diffusion coefficients (Fig. 9) after EuTc exposure when compared to the recovery rates and diffusion coefficients of DOPC/TR-DHPE and DOPC/TopFluorPC. The DOPC/BODIPY-GM1 diffusion coefficient drops from 1.78 ± 0.25 μm^2^/s to 0.54 ± 0.07 μm^2^/s after the addition of EuTc. The fact that the diffusion coefficients after EuTc are comparable to those with TR-DHPE and TopFluorPC suggests that regardless of the probe employed, EuTc significantly reduces lipid diffusion. Interestingly, Kutsenko and coworkers found that β-diketone complexes of Eu^3+^ caused alterations of bilayer membrane properties.^41^ Of particular relevance is their finding that these complexes induce tighter packing in the hydrophobic region of the bilayer, which may help explain the reduction in lipid diffusion coefficients that we observe.

An additional series of FRAP experiments was conducted to determine if Eu^3+^ or Tc can independently modulate the diffusion of fluorescent lipid probes in SLBs. In these experiments SLBs possessing one of a variety of fluorescent probes (TR-DHPE, TopFluorPC, BODIPY-GM1, or TopFluor-cholesterol) were examined by FRAP before and after exposure to 1 μM Eu^3+^ or 1 μM Tc in MOPS buffer. These experiments showed that both the Eu^3+^ and Tc had no or little impact on lipid diffusion coefficients (Fig. S9), which suggests that it is the properties of the EuTc complex, rather than the central Eu^3+^ or the Tc ligand that causes reduced lipid mobility. Because EuTc decreases lipid diffusion coefficients of a range of fluorescent probes bearing different chemical functionality, it is unlikely that EuTc specifically interacts with fluorescent probes, but rather interacts primarily with the background PC lipids in the bilayer, which may be in part responsible for redistribution of GM1 observed in GUV imaging experiments.

### Cellular membrane labeling with EuTc

To determine if EuTc would be useful for illuminating membranes in a cellular context, we used EuTc to label UMSCC-2 cells. This cell line originates from a head and neck squamous cell carcinoma.^42^ The UMSCC-2 cells were seeded on to poly-L-lysine coated coverslips, then incubated with 10 μM EuTc in MOPS buffer for 20 minutes. After the incubation period, the cells were imaged in brightfield, then their EuTc luminescence was examined (Fig. 10A-B).

**Figure 10.**
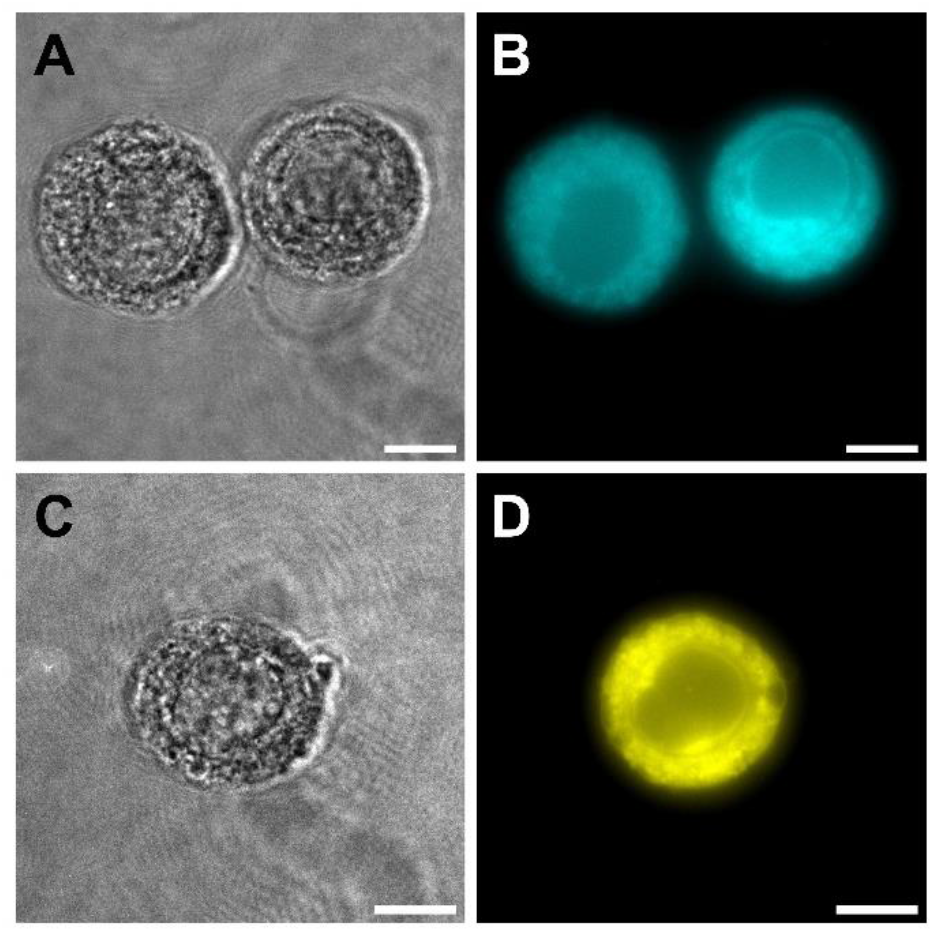
Cellular membrane labeling with EuTc. (A-B) UMSCC-2 cells incubated with EuTc and then imaged in brightfield (A) and EuTc (B) channels. (C-D) A UMSCC-2 cell incubated with FM1-43 then imaged in brightfield (C) and FM1-43 (D) channels. All scale bars are 10 μM.

To compare the EuTc labeling of UMSCC-2 cells to that of a conventional membrane stain, we treated cells with 4 μM FM1-43 in MOPS, and imaged cells in brightfield and fluorescence (Fig. 10C-D). FM1-43 is a water soluble stain that is nonfluorescent in aqueous solutions. However, FM1-43 can partition into lipid bilayer membrane, whereupon its fluorescence emission increases manyfold.^43^ In comparing the EuTc and FM1-43 fluorescence images, we observe similar staining patterns, indicating that EuTc could potentially find utility in cellular imaging applications where its large Stokes shift and extremely narrow emission band are beneficial. While the toxicity of EuTc toward UMSCC-2 cells was not examined, it is interesting to note that Eu^3+^ (administered as EuCl_3_) has low toxicity in mice. Haley and coworkers measured an acute intraperitoneal LD_50_ of 550 mg/kg and an acute oral LD_50_ of 5000 mg/kg.^44^

## CONCLUSIONS

In conclusion, this work demonstrates the application of a coordination complex probe suitable for imaging spatial heterogeneity of biomembranes. Compared to traditional organic probes for biomembranes, EuTc has a much narrower emission bandwidth, an extremely large Stokes shift, and a unique combination of excitation and emission wavelengths. Additionally, EuTc has a much longer emission lifetime (tens of μs)^20^ than most organic membrane probes. These attractive properties suggest that EuTc may find applications in multicolor fluorescence imaging and time-resolved imaging of biomembranes, with the caveat that some lipid redistribution may occur. In particular, the unique spectral features of EuTc could help reduce or eliminate fluorescence crosstalk between imaging channels in multicolor fluorescence imaging applications. We show that introducing EuTc to liposome suspensions results in a lipid concentration-dependent increase in EuTc luminescence. Additionally, when labeling phase separated GUVs with the EuTc complex, we observe a distinct contrast between Lo and Ld domains, where the EuTc labels the Ld domains. Surprisingly, labeling GM1-containing, phase-separating GUVs with EuTc results in GM1 losing its preference for the Lo phase. In addition, we observe significant shifts to lipid probe diffusion coefficients after EuTc is added to membranes, which suggest nonspecific changes to membrane lipid order.

Of course, it is not ideal for a membrane label to cause spatial redistribution of the lipids. At this time it is unclear whether it is only the gangliosides that are being redistributed, or if other lipids are redistributed as well. If redistribution only applies to gangliosides, then EuTc would be useful in labeling ganglioside-free GUVs, especially in situations where multicolor imaging is difficult with traditional organic probes due to crowded spectral space. Our experiments with GUVs displaying Ld domains with predictable area fractions (Fig. 5) indicate that EuTc may not significantly alter the distribution PCs and cholesterol, at least as far as it is detectable with the TR-DHPE probe. While beyond the scope of the present investigation, it would be possible to determine whether EuTc causes cholesterol redistribution using fluorescent analogs of cholesterol, for example TopFluor-cholesterol, which has been shown to selectively partition into Lo domains.

## METHODS

### Reagents and Chemicals

Dioleoylphosphatidylcholine (DOPC), dipalmitoylphosphatidylcholine (DPPC), ganglioside GM1 (ovine brain), 1-palmitoyl-2-(dipyrrometheneboron difluoride) undecanoyl-sn-glycero-3-phosphocholine (TopFluorPC), TopFluor-cholesterol, and cholesterol were all purchased from Avanti Polar Lipids (Alabaster, AL). Sucrose, chloroform, europium (III) chloride hexahydrate (99.99%), tetracycline hydrochloride (cell culture grade, >95%), and cholera toxin B subunit FITC conjugate (CTxB-FITC) were purchased from Sigma Aldrich. 3-(N-morpholino)propanesulfonic acid) (MOPS) was purchased from Acros Organics. Texas Red DHPE (TR-DHPE), BODIPY FL C_5_-ganglioside GM1 (GM1-BODIPY) (Invitrogen), cholera toxin B subunit Alexa Fluor 647 tagged (CTxB-Alexa647), and sodium dodecyl sulfate (SDS) were purchased from ThermoFisher.

### EuTc Preparation

In all experiments, EuTc was prepared by combining equimolar amounts of europium (III) chloride and tetracycline hydrochloride in 10 mM MOPS, pH 7.0. Prior to their combination, europium (III) chloride and tetracycline hydrochloride were kept as stock solutions in 10 mM MOPS, pH 7.0. The tetracycline hydrochloride stock solution was prepared fresh daily. The EuTc solutions, prepared fresh daily, typically contained 1 mM europium (III) chloride and 1 mM tetracycline hydrochloride. Unless otherwise stated, the EuTc solution was then diluted to 1 μM for spectroscopy and imaging. The EuTc complex was added to liposome and GUV samples after their preparation.

### Liposome Preparation

Lipids dissolved in chloroform were mixed in glass vials to their desired molar ratios with a final concentration of 1.00 mg/mL. Chloroform was evaporated under vacuum at room temperature for a minimum of 2 h. Lipid films were rehydrated in MOPS buffer (10 mM MOPS, pH 7.00), vortexed and subjected to three freeze-thaw cycles. Freezing was accomplished by plunging the sample into liquid N_2_ until frozen, and then samples were thawed with a warm water bath. Liposomes were then extruded inside a mini-extruder (Avanti) using a 100 nm pore size polycarbonate membrane filter (Whatman) for a total of 23 passes. Immediately after, liposomes were used in fluorescence spectroscopy experiments and to form supported lipid bilayers (SLBs).

### Preparation of GUVs

Giant unilamellar vesicles (GUVs) were made by electroformation.[32] Lipid mixtures were dried under vacuum, resuspended in chloroform, and then a droplet of lipid solution was transferred to clean indium tin oxide (ITO) coated slides (Delta Technologies, Loveland, CO). ITO slides were cleaned by swabbing the slides with 2% aqueous Alconox detergent solution and rinsing with MilliQ H_2_O immediately thereafter, three times over. Lipid solutions on the ITO slides were dried under vacuum for a minimum of 15 min to remove chloroform. A capacitor was formed with a second ITO coated slide with a 0.3 mm Teflon spacer. The two slides were sandwiched together using binder clips. 400 µL of 200 mM sucrose was injected between the two slides. The total lipid concentration in the chamber was 1.0 mg/mL. GUVs were electroformed at 65 °C, using an AC signal with peak to peak amplitude of 3.0 V and frequency 10 Hz for 2 h (Siglent Technologies). GUVs were removed from the electroformation chamber and diluted 1:1000 with MOPS buffer in an Attofluor imaging chamber (ThermoFisher Scientific). Prior to imaging, GUVs were exposed to 1 µM EuTc and/or 10 nM CTxB-Alexa647. A minimum of 30 GUVs among ≥ 3 individual preparations were examined for signal partitioning and representative images were chosen to produce in the figures.

### Fluorescence Spectroscopy

Excitation and emission spectra were collected with a Fluorolog-2 spectrofluorometer (Horiba). Excitation spectra for EuTc were collected by scanning excitation from 350-450 nm while monitoring emission at 618 nm. EuTc emission spectra were collected from 560-700 nm by exciting at 400 nm. TR-DHPE excitation was collected by scanning from 450-605 nm while monitoring emission at 615 nm. Emission spectra for TR-DHPE were collected by scanning from 600-650 nm while exciting at 596 nm. EuTc (1 µM) emission was measured in the presence of DOPC liposomes (0.01 µM-100 µM total lipid concentration). TR-DHPE emission was measured from 1 µM DOPC liposomes containing 1 mol % TR-DHPE.

### Fluorescence Microscopy of GUVs

Glass coverslips were cleaned with 2 % (w/v) SDS solution and rinsed with ultrapure H_2_O, and then dried with N_2_ gas. Clean glass coverslips then underwent a UV-ozone treatment (UV/Ozone ProCleaner Plus, BioForce Nanosciences) for approximately 10 min. To label GUVs with EuTc and CTxB-Alexa647, they were first exposed to 1 µM EuTc, incubated for 5 min, then exposed to 10 nM CTxB-Alexa647 (or vice-versa), incubated for 5 min, and imaged immediately after. All imaging was done using an inverted microscope (Eclipse Ti, Nikon) equipped with a 100× oil immersion objective with a 1.49 numerical aperture. Fluorescence was excited using a LED light engine (Aura II, Lumencor), a Cy5 (Chroma), FITC (Chroma), TRITC (Chroma) or a custom EuTc (Semrock) filter set. All images were captured with a 2048 × 2048 pixel sCMOS camera (Orca Flash 4.0 v2, Hamamatsu) controlled by Nikon Elements software.

### Supported Lipid Bilayer Preparation for Fluorescence Recovery after Photobleaching (FRAP)

Glass coverslips were cleaned with 2% (w/v) SDS and rinsed with ultrapure H2O and then dried with N2 gas. The coverslips were subjected to a 10 min UV-ozone treatment and adhered to a self-adhesive bottomless 6-channel slide (Sticky-Slide VI 0.4, Ibidi) or mounted in Attofluor chambers. Vesicles to form SLBs contained DOPC and 1 mol % TopFlourPC, 1 mol % TR-DHPE, 1 mol % TopFluor-cholesterol, or 1 mol % BODIPY-GM1. The liposomes were diluted to 0.1 mg/mL in Tris buffer (10 mM Tris, 150 mM NaCl, pH = 7.0) and then incubated on the coverslips for 30 min. After incubation, excess liposomes were first washed out with Tris buffer, and then the buffer was exchanged for MOPS (pH = 7.0). After the SLBs were washed, they were imaged with an inverted microscope (Eclipse Ti, Nikon) equipped with a 100× oil immersion objective with a 1.49 numerical aperture. Fluorescence was excited using a LED light engine (Aura II, Lumencor), a FITC or TRITC filter set (Chroma) and images were captured with 2048 × 2048 pixel sCMOS camera (Orca Flash 4.0 v2, Hamamatsu). Each sample was photobleached using a 405 nm laser (50 mW) pulse for 2 s, and fluorescence recovery was captured at 1 s intervals for 60-90 s. After FRAP, the samples were exposed to 1 µM EuTc, 1 μM EuCl_3_, or 1 μM Tc, incubated for 1 h and photobleached again. Recovery was monitored and recorded for 1 s intervals for 60-90 s and captured using a TRITC or FITC filter set. Lipid diffusion coefficients were calculated using the Hankel transformation and MATLAB code described by Jonsson et al.^45^

## Author Contributions

J.L.C., A.I.M., A.R.H-S., and N.J.W. designed experiments, J.L.C., B.A.B., A.T.O., D.E.S., and A.N.S. carried out the experiments; J.L.C., B.A.B., A.T.O, D.E.S, A.N.S., A.R.H-S., and N.J.W. analyzed the data; J.L.C. and N.J.W. wrote the manuscript with input from all co-authors. B.A.B., A.T.O., D.E.S., and A.N.S. contributed equally.

## Acknowledgements

N.J.W. acknowledges funding from Lehigh University and the National Science Foundation (Award Number 2044792). The authors thank Damien Thévenin and Eden Sikorski for providing the UMSCC-2 cells.

## Supplementary Information

Supplementary material contains custom filter cube characteristics, additional images of GUVs labeled with EuTc, TR-DHPE, and CTxB-FITC, a lipid composition phase diagram, fluorescence intensity profiles for GUVs, and a summary of additional FRAP data, including fluorescence images.

## For Table of Contents Only

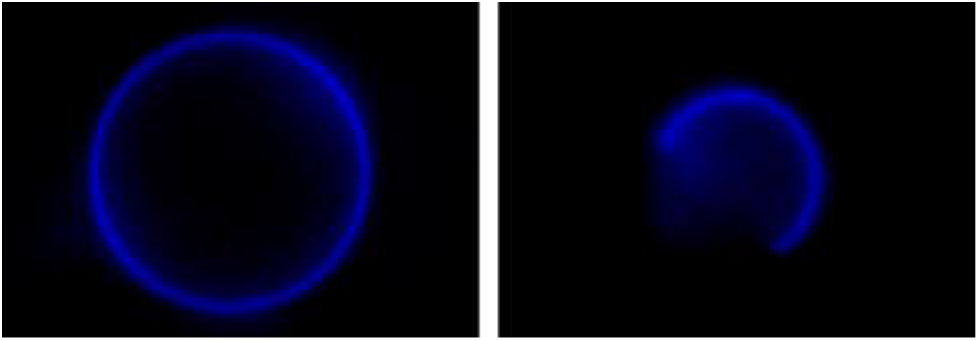

