## Supplemental information for "Imaging Giant Vesicle Membrane Domains with a Luminescent Europium Tetracycline Complex"

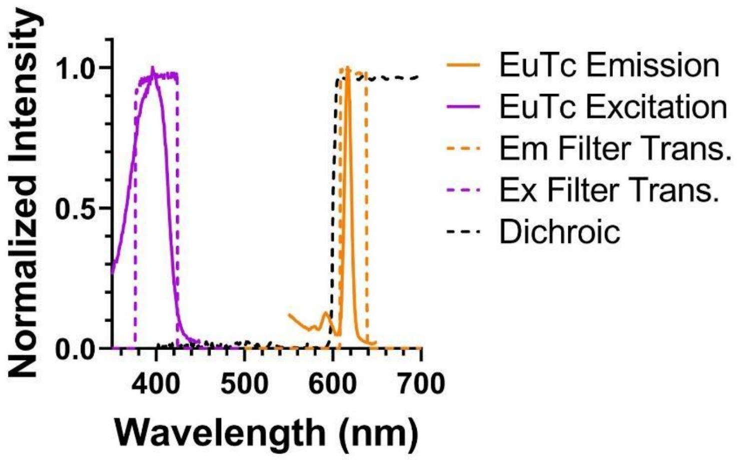


**Figure S1.** Characteristics of the custom filter cube for microscopic imaging of EuTc superimposed over the excitation and emission spectra of EuTc. The dashed lines indicate the transmission of the excitation, dichroic, and emission filters.

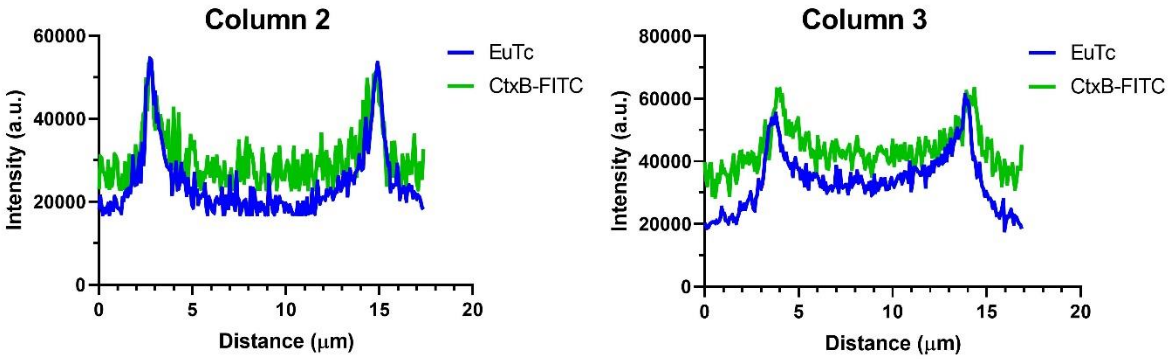


**Figure S2.** Fluorescence intensity profiles taken along the lines shown in Figure 4 of the main text. The left and right figures correspond to Columns 2 and 3, respectively. The lipid compositions are as follows: Column 2: DOPC/GM1 (98:2); Column 3: DOPC/GM1/cholesterol (78:2:20).

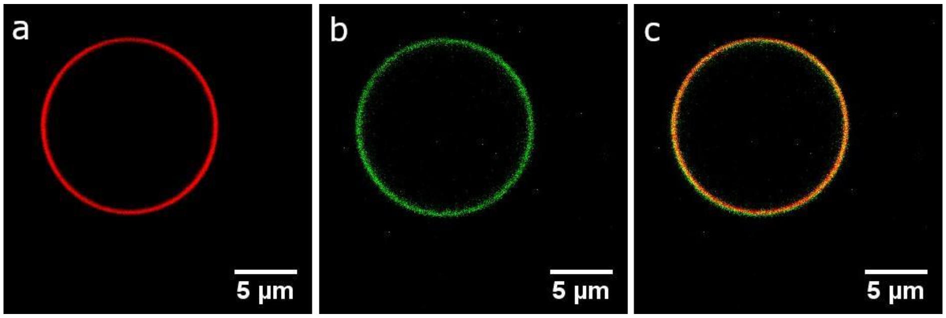


**Figure S3.** DOPC/GM1/TR-DHPE (99:1:1) GUV imaged by (a) TR-DHPE fluorescence and (b) CTxB-FITC fluorescence. (c) Merged images of (a) and (b) result in complete colocalization of CTxB-FITC and TR-DHPE fluorescence.

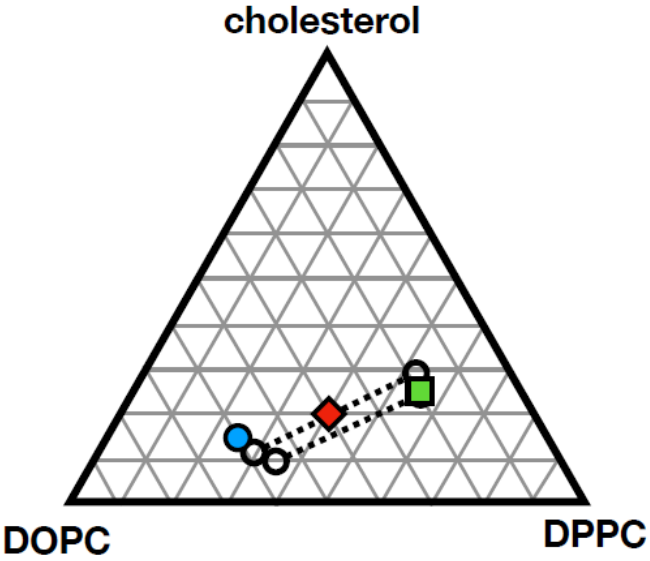


**Figure S4.** Phase diagram showing the compositions of vesicles used in the three columns of Figure 5 of the main text. The GUVs were composed of DOPC, DPPC, cholesterol with varying ratios, and all vesicles contained 1% TR-DHPE. The compositions are as follows (DOPC:DPPC:cholesterol): Column 1 (square), 19:55:25; Column 2 (diamond), 39.5:39.5:20; and Column 3 (filled circle), 59:25:15. These compositions are superimposed on the approximate location of two local tie lines in the liquid-liquid coexistence region (dotted lines with open symbol endpoints) measured quantitatively at 25° C by fitting NMR spectra for the same ternary mixture of lipids.[1]

When two phases coexist, the chemical composition of each phase can be identified on the phase diagram from the endpoints of the "tie-line" passing through the point which represents the overall composition.[2]

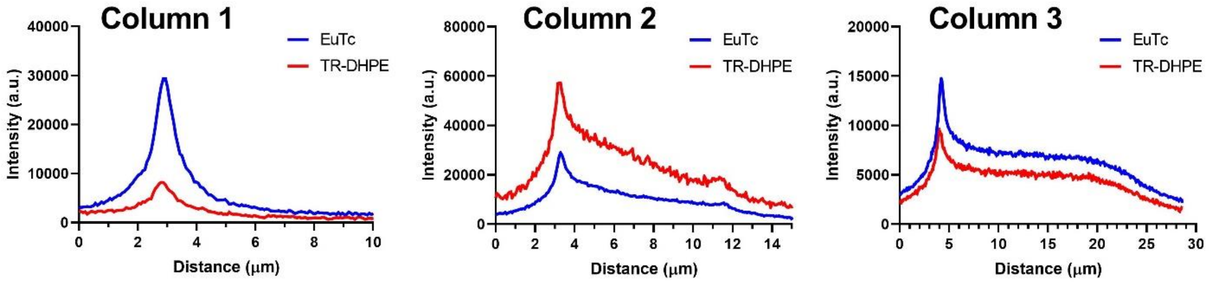


**Figure. S5.** Fluorescence intensity profiles taken along the lines shown in Figure 5 of the main text. The left, center, and right figures correspond to the intensity profiles of the lines shown in Column 1, Column 2, and Column 3 of Figure 5, respectively. The compositions of the GUVs are as follows: Column 1: DOPC/DPPC/cholesterol/TR-DHPE (19:55:25:1); Column 2: DOPC/DPPC/cholesterol/TR-DHPE (39.5:39.5:20:1); Column 3: DOPC/DPPC/cholesterol/TR-DHPE (59:25:15:1)

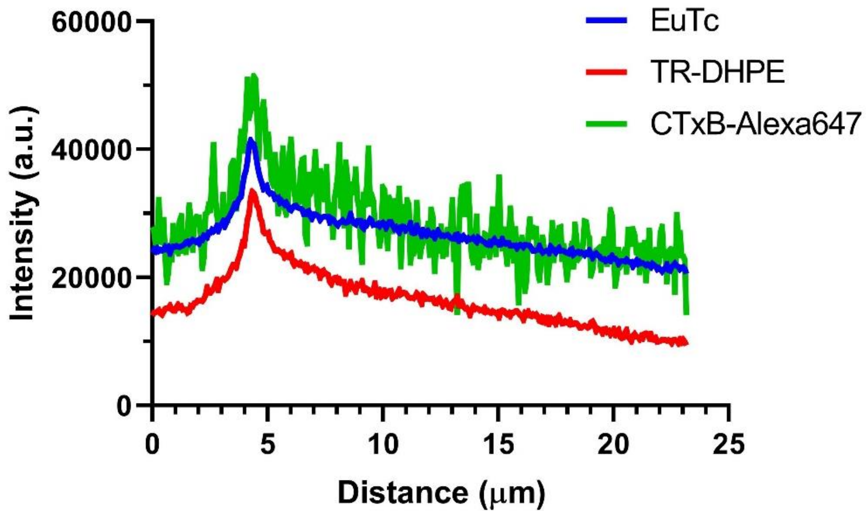


**Figure S6.** Fluorescence intensity profiles along the dashed line in Figure 6, column 3 of the main text.

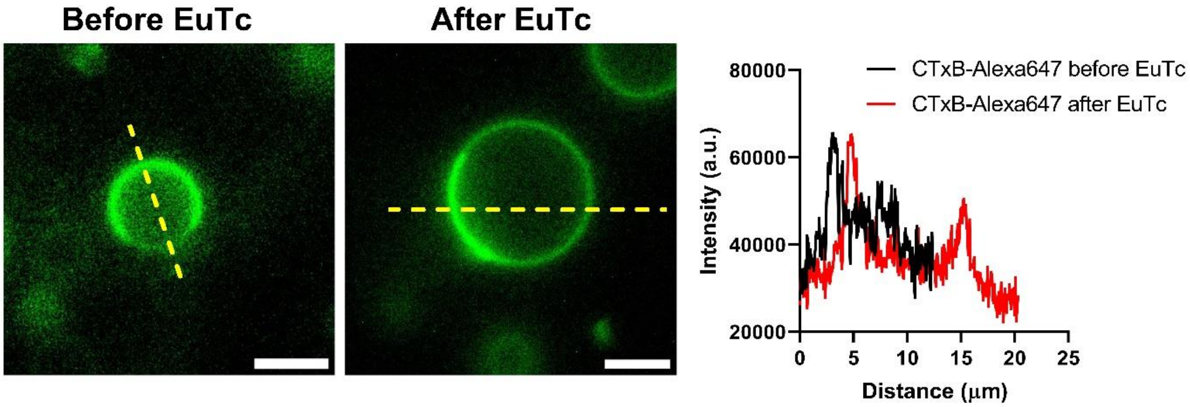


**Figure S7.** Fluorescence images and intensity profiles for CTxB-Alexa647 staining of GUVs in the absence and presence of EuTc. The GUV was composed of DOPC, DPPC, cholesterol, and GM1 with the following molar ratio: 39.5:39.5:20:1.

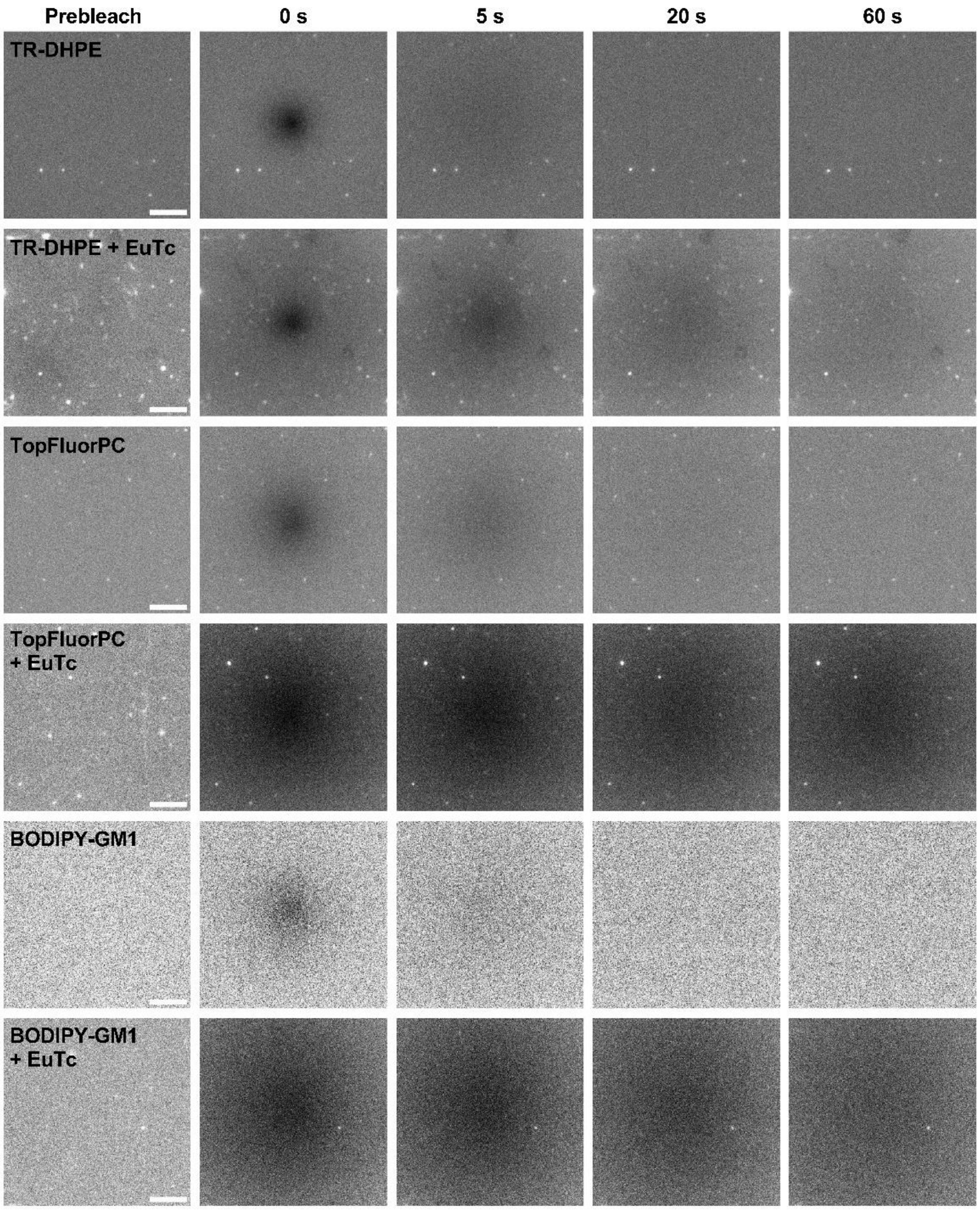


**Figure S8.** Fluorescence micrographs from representative FRAP experiments. The recovery curves summarizing these FRAP experiments are shown in Fig. 8 in the main text.

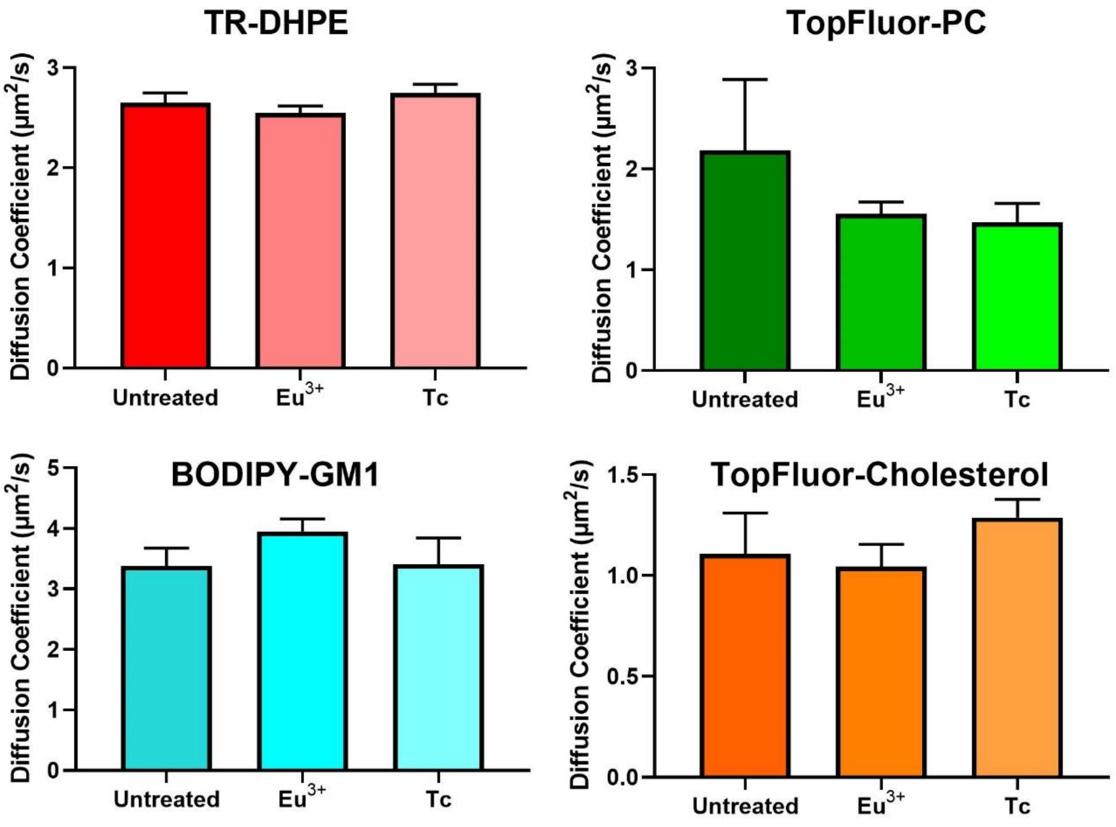


**Figure S9.** Diffusion coefficients of TR-DHPE, TopFluor-PC, BODIPY-GM1, and TopFluor-Cholesterol in POPC SPBs measured by FRAP before and after exposure to 1 μM $Eu^{3+}$ or 1 μM Tc in MOPS buffer, pH=7.0.

### Supporting Information References

1. Veatch, S. L.; Soubias, O.; Keller, S. L.; Gawrisch, K., Critical fluctuations in domain-forming lipid mixtures. *Proc Natl Acad Sci U S A* **2007,** *104* (45), 17650-5.
2. Ferguson, F. D.; Jones, T. K., *The phase rule*. Butterworths: London, **1966**.
